# Functional ultrasound imaging reveals an aberrant developmental trajectory of functional connectivity in a mouse model of the 15q13.3 copy number variant microdeletion

**DOI:** 10.1101/2022.10.24.513480

**Authors:** Gillian Grohs-Metz, Bastian Hengerer, Hugo Cruces Solis

## Abstract

The developmental nature of many neuropsychiatric disorders such as schizophrenia necessitates the detection of functional biomarkers during the prodromal phase of disease that can predict symptomatic conversion and outcomes. Structural chromosomal aberrations, such as copy number variants, confer high risk and penetrance of neuropsychiatric disorders. We used functional ultrasound imaging (fUS) to characterize the functional connectivity profile of the 15q13.3^+/−^ copy number variant mouse model during major developmental milestones on post-natal day (p)35, 42, 60, and 90 in comparison to wild type littermates. We identified divergent trajectories for 15q13.3^+/−^ mice and WT littermate controls where functional connectivity was reduced for both genotypes with age, but to a lesser extent for 15q13.3^+/−^ mice. We were then able to isolate the distinct differences between genotypes to identify a large-scale network where 15q13.3^+/−^ mice displayed global cortical hyperconnectivity and elevated intra-connectivity within the hippocampus and amygdala, in particular. In order to determine the stage of development where the connectivity trajectories bifurcated, we used machine learning to predict genotype. We found that the connectivity profile from p42, but not p35, predicted the genotype of individual mice at p90 with 82% accuracy. All together, these results suggest a crucial period of network maturation from early to late pubescence that is pivotal in the transition of healthy network connectivity into adulthood. This novel application of fUS longitudinally through development shows promise in improving the understanding of the disease biology of mouse models of psychiatric diseases.

## Introduction

The ability to detect and measure functional biomarkers in developmental models offers great promise to guide early intervention in psychiatric diseases such as schizophrenia. Frequently defined as a neurodevelopmental disorder, the first episode of psychosis typically presents during late adolescence to early adulthood between the ages of 15 to 30 (Jones, 2013). However, there is substantial evidence that physiological deviations in neural networks are evident prior to the manifestation of pronounced symptoms during the prodromal phase of disease (Allen et al., 2012; Wolf et al., 2015). As an organism develops, the transition from overall excitatory to inhibitory control of neural activity is accomplished through cellular processes such as synaptic pruning, myelination and increased inhibitory signaling. These processes effectively refine and strengthen the integration between various networks to improve signal processing efficiency (Marek et al., 2015).

Functional connectivity, a measure of network integration through the temporal correlation of regional neural activity, has emerged as a highly informative approach to understanding the basis of cognition, behavior, and mood. As neural networks are considered to be one of the foundations of brain function, it is unsurprising that virtually all psychiatric disorders display at least some circuit aberrations, where differences range from hypo- to hyperconnectivity and generally associate with symptom severity (McTeague et al., 2017, 2020). Dysfunction in executive control circuitry, abnormal threat responses in the amygdala, and differences in hippocampal connectivity have been detected in patients with schizophrenia and unaffected family members at high risk of developing psychosis (Allen et al., 2012; Wolf et al., 2015; Edmiston et al., 2020). In addition, abnormalities in sensorimotor processing and integration are particularly prevalent, have been found to correlate with age of onset, and are thought to contribute significantly to the profound positive symptoms of schizophrenia (Manschreck et al., 2004, 2015; Minassian et al., 2007; Carment et al., 2019). Furthermore, deviations in the organization of functional network connectivity in youth have been shown to reliably predict the conversion to psychosis (Collin et al., 2020b, 2020a). Pinpointing specific changes in the transition from risk to the presentation of psychosis will enable strategic therapeutic targeting that could drastically improve patient quality of life and prevent the substantial health, social, and economic burdens associated with mental illness.

Disorders with chromosomal abnormalities which encompass several genes, such as copy number variants (CNVs), are prime candidates for preclinical disease modeling. While these variants are relatively rare in the general population, they confer a very high risk of developing psychiatric disorders such as schizophrenia (Odds ratio = 2 - 57) (Rees et al., 2014), an attribute that enables the study of the developmental processes leading to schizophrenia. One such CNV consisting of a microdeletion in the 15q13.3 locus of chromosome 15, has been shown to confer a high risk of developing schizophrenia (Odds ratio = ~7.5), as well as autism, intellectual disability, and epilepsy (Rees et al., 2014). A mouse model of this syndrome has been previously developed and expresses many of the features of the human 15q13.3 microdeletion syndrome. Similar to that seen in schizophrenia sufferers, these animals display auditory processing deficits, an increase in gamma oscillations at rest but a decrease after an auditory stimulus, aggressive behaviors after stress, impaired spatial reference memory, and also develop increased body weight (Fejgin et al., 2014). Furthermore, these mice exhibit hyperconnectivity throughout multiple brain regions (Gass et al., 2016), which is frequently correlated with multiple aspects of schizophrenia (Klingner et al., 2014; Krishnadas et al., 2014). It should be noted, however, that these preclinical studies were conducted in adult mice. As alluded to previously, understanding and detecting the subtle changes throughout development that lead to this phenotype, are crucial in not only developing more effective therapies, but in finding precise biomarkers to help identify and classify patients at risk for developing schizophrenia.

The novel functional ultrasound (fUS) imaging modality provides a valuable tool to investigate neural circuitry at the whole brain scale. The non-invasive nature of fUS imaging enables network wide coverage with an exceptionally high signal-to-noise ratio (Macé et al., 2011; Osmanski et al., 2014; Gesnik et al., 2017). fUS employs ultrafast, plane wave technology to measure Doppler signal from blood flow. It has a higher resolution than many other imaging modalities such as positron emission tomography and many MRI machines, and is less restraining in terms of magnetic objects, noise, mechanical ventilation, and physical access to animals (Catana, 2019; Mandino et al., 2020; Deffieux et al., 2021). These combined attributes render fUS imaging an ideal technique for longitudinal studies of neural networks in mice in the context of psychiatric disease. Here, we profiled functional connectivity in a mouse model of 15q13.3 CNV during developmental milestones that reflect transitions in neural network development in humans. Furthermore, we aimed to isolate a critical period of reorganization in functional connectivity that could predict the established neural network signature at adulthood.

## Materials and methods

### Animals

All experiments were approved by the appropriate authority (Regierungspräsidium Tübingen, Germany) in accordance with European Directive 2010/63/EU and performed in an AAALAC (Association for Assessment and Accreditation of Laboratory Animal Care International) accredited facility. Male Df(h15q13)^+/−^ mice and Df(h15q13)^+/+^ littermates, hereafter referred to as 15q13.3^+/−^ and WT, were generated as previously described (Fejgin et al., 2014) by TaconicArtemis (Köln, Germany) and bred and shipped by Lundbeck (Valby, Denmark). Animals were housed in groups of 4 on a normal 12:12 h light cycle and habituated for 7 days before any experiments were performed. Imaging was performed on post-natal day 35 (p35), p42, p60, and p90 (Fig.1A).

### fUS image acquisition and data pre-processing

Animals were lightly sedated with dexmedetomidine similar to previously described methods (Nasrallah et al., 2014; Boido et al., 2019; Grohs-Metz et al., 2022). Briefly, animals were initially anesthetized with 4 % isoflurane and given a subcutaneous analgesic for any pain that might be experienced from ear bar fixation (Metacam, Boehringer-Ingelheim, 0.5 mg kg^−1^). Once animals were fixed in a stereotactic frame (David Kopf Instruments, USA) positioned on an anti-vibration table (CleanBench™, TMC, Germany), isoflurane was reduced to 1 % and temperature was maintained at 37 °C with a homeothermic warming system. Sedation was maintained with a subcutaneous 0.067 mg kg^−1^ bolus of dexmedetomidine (Tocris, USA; freshly prepared and filtered), followed by a continuous infusion of 0.20 mg kg^−1^ hr^−1^ at a flow rate of 5 mL kg^−1^. After 10 minutes of dexmedetomidine infusion, isoflurane was gradually reduced by 1 % per minute until full cessation 20 minutes after initial bolus. Imaging was not initiated until 15 minutes after cessation of isoflurane.

Hair was closely shaved from the scalp and an isotonic coupling gel applied between the probe and the skin. A dedicated small animal fUS device was used to acquire and process images (Iconeus, France) as previously described in depth(Macé et al., 2011; Osmanski et al., 2014; Demené et al., 2015; Gesnik et al., 2017). In short, a 128 piezoelectric, linear, ultrasonic probe with a central frequency of 15 MHz connected to an ultrafast ultrasound scanner resulted in a final sequence of 2.5 Hz, 100 x 100 x 400 μm power Doppler functional images. An integrated singular value decomposition filter enabled subtraction of tissue noise to extract pure blood signal (Demené et al., 2015). A series of 2-D images were used to register that brain in real time using the vascular Iconeus Brain Positioning System (Nouhoum et al., 2021) (Fig. 1B). Images were then acquired in the plane of interest for 10 minutes to measure resting state connectivity. After imaging, dexmedetomidine sedation was rapidly reversed with Atipamezole (Alzane, Zoetis, 5 times total dexmedetomidine dose) for full recovery.

**Fig. 1.**
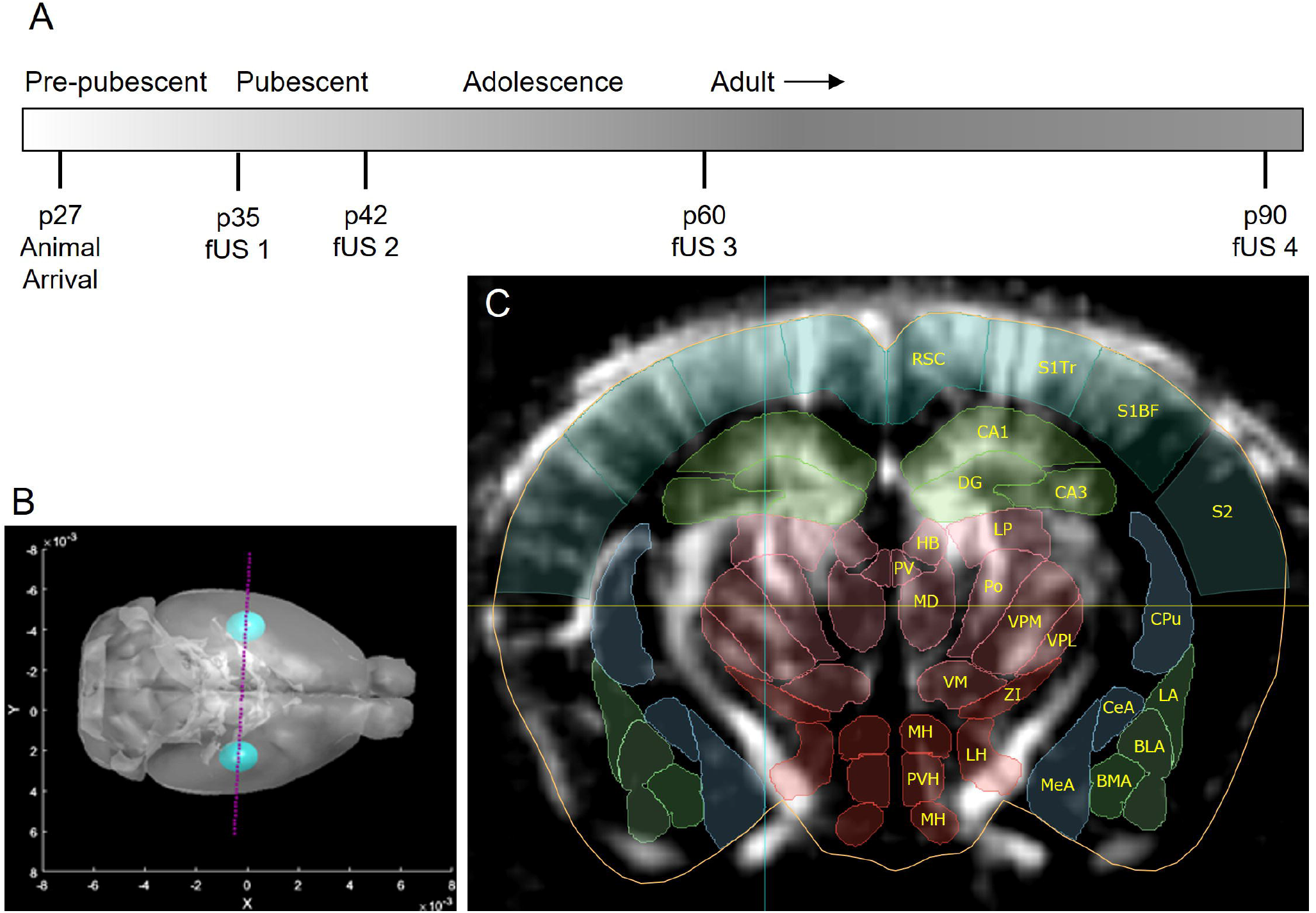
Experimental schedule and fUS imaging plane. (**A**) Animals were imaged during early and late pubescence, the transition to adulthood, and again during full adulthood. (**B**) During each fUS imaging session, the brain was registered in 3D to enable precise probe placement (purple dotted line **B**) for coverage of cortical, hippocampal, thalamic, amygdalar, and hypothalamic regions of interest (ROIs) (**C**). *RSC* retrosplenial area, *S1BF/S1Tr* primary somatosensory area, barrel field/trunk, *S2* supplementary somatosensory area, *CA1/CA3* Ammon’s horn field CA1/CA3, *DG* dentate gyrus, *LP/MD/PV* lateral posterior/mediodorsal/ periventricular nucleus of the thalamus, *HB* Habenula, *Po* posterior complex of the thalamus, *VM/VPL/VPM* ventral medial/ventral posterolateral/ventral posteromedial nucleus of the thalamus, *ZI* Zona incerta, *LH/MH/PVH* lateral/medial/periventricular hypothalamic area, *CPu* caudoputamen, *BLA/BMA/LA* basolateral/basomedial/lateral amygdalar nucleus, *CeA/MeA* central/medial amygdalar nucleus

### Region of interest selection and functional connectivity calculation

Individual animal 3D brain registrations were used post hoc to delineate anatomical ROIs using the Allen Mouse Brain Common Coordinate Framework (CCFv3) (Wang et al., 2020) (Fig. 1C). Temporal power Doppler signal (10 min) was extracted from each voxel within an ROI to calculate an average cerebral blood flow (CBV) signal for each ROI. The signal was normalized using a 4^th^ order polynomial fit function and a 0.2 Hz low pass filter was used to isolate frequencies of functional connectivity (Macé et al., 2011; Nasrallah et al., 2014; Osmanski et al., 2014). Correlation coefficients were calculated for each ROI-ROI pair using Pearson product moment correlation which were normalized using Fisher’s *z* transformation to create connectivity matrices for individual animals for each imaging session.

### Data analysis

To improve the visualization of the correlation matrices, we used correlation matrix-based hierarchical clustering to group brain regions with similar connectivity patterns (Fig. 2). To analyze developmental changes in functional connectivity, we calculated the effect size (Cohens’ *d*) using the correlation values as previously described (Zerbi et al., 2019) and used the WT correlation values at p35 as reference. To obtain a measure of change in connectivity per developmental time point, we averaged all the effect size values of each brain region and then calculated the mean across all brain regions.

**Fig. 2.**
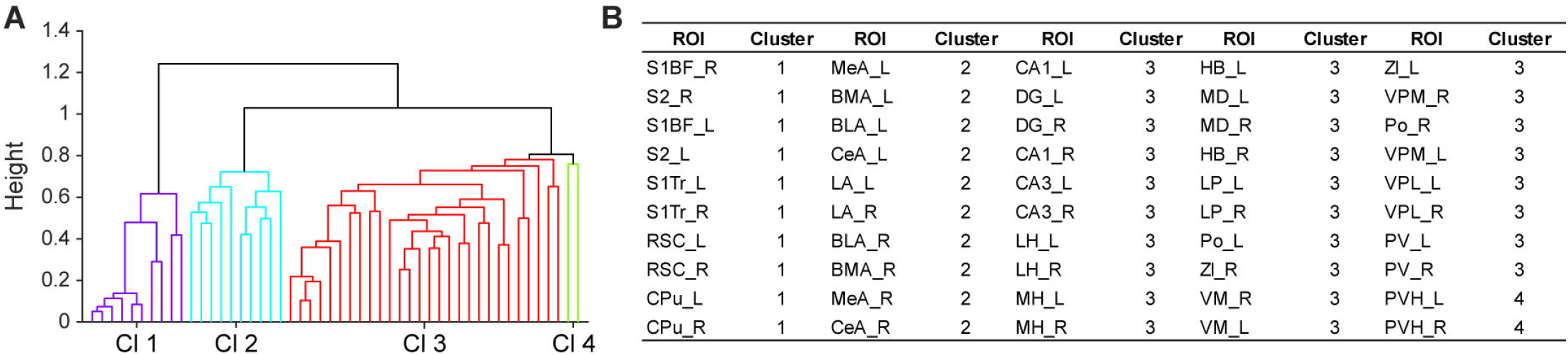
Functional connectivity-based ROI clustering. (A) Cluster hierarchy. (B) Cluster designation. Cluster 1 contains mainly cortical ROIs and the CPu. Cluster 2 contains the ROIs of the amygdalar nuclei. Cluster 3 contains hippocampal, thalamic subdivisions, and a majority of the hypothalamic nuclei. Cluster 4 contains the bilateral paraventricular hypothalamic nuclei. L and R designators signify left and right hemispheric regions, respectively. ROI abbreviations are defined in Fig. 1.

To identify changes in functional connectivity between genotypes, we used the network-based statistics toolbox (NBS) in MATLAB, a validated method to perform statistical analysis on large networks (Zalesky et al., 2010). For the input data, we used the original *z* transformed correlation values of the entire dataset and used a repeated measures design with a primary threshold of 3 and 5000 permutations. The resulting t-values (edges) and numbers of significant connections of a particular ROI (node degree) were used to construct and visualize a distinct network of hyperconnectivity in the 15q13.3^+/−^ mice.

To predict the genotypes based on changes in functional connectivity, we first performed a principal component analysis (PCA) using the Cohen’s *d* effect size values of the entire dataset (*pca*, function). Next, we used the scores of the first three principal components as predictors to train a linear support vector machine (SVM) (*fitcsvm*, function). To protect against overfitting, we partitioned the data into 5-fold cross validation (*crossval*, function). To predict the genotypes at p90 with data from p35 or p42, we used the same method described above but we calculated independent PCA scores for each age. In this way, the data used for training and predicting were independent of each other.

## Results

To characterize functional connectivity across developmental stages, we longitudinally imaged the same mice across different time points (p35, p42, p60 and p90), thereby monitoring the transitions from pubescence to adolescence and adulthood. Some animals awakened during fUS imaging, therefore only animals that were stable for all 4 imaging sessions were analyzed (WT n=11, 15q13.3^+/−^ n=12). To identify the impact of age on functional connectivity, we computed the effect size using the correlation values of WT mice at p35 as reference (Zerbi et al., 2019). In WT animals we observed a decrease in functional connectivity with increasing age (Fig. 3A, top row cool colors). This effect was accurately reflected when we computed the average effect size for each developmental stage (Fig. 3B, black bars). In contrast, 15q13.3^+/−^ mice showed a distinct pattern of functional connectivity at p35 (Fig. 3A, bottom row) where most of the functional connectivity values were lower than those of WT at p35 (Fig. 3B, red bars). At p42, the changes in functional connectivity were similar between genotypes. At p60, we found that the pattern of functional connectivity in 15q13.3^+/−^ mice was different than in WT for a subset of brain regions. Mainly, regions in cluster 1 seemed to have higher functional connectivity with regions in other clusters, while regions in cluster 2 showed higher intra-functional connectivity (Fig. 3A, third column). However, the average change in functional connectivity between genotypes was not different (Fig. 3B). At p90, the values of functional connectivity of 15q13.3^+/−^ mice did not decrease as in WT and were similar to those observed at p60 (Fig. 3A, fourth column and Fig. 3B). Together, these data indicate that functional connectivity in 15q13.3^+/−^ mice is already divergent at p35 and follows a different developmental trajectory that results in hyperconnectivity in adulthood.

**Fig. 3.**
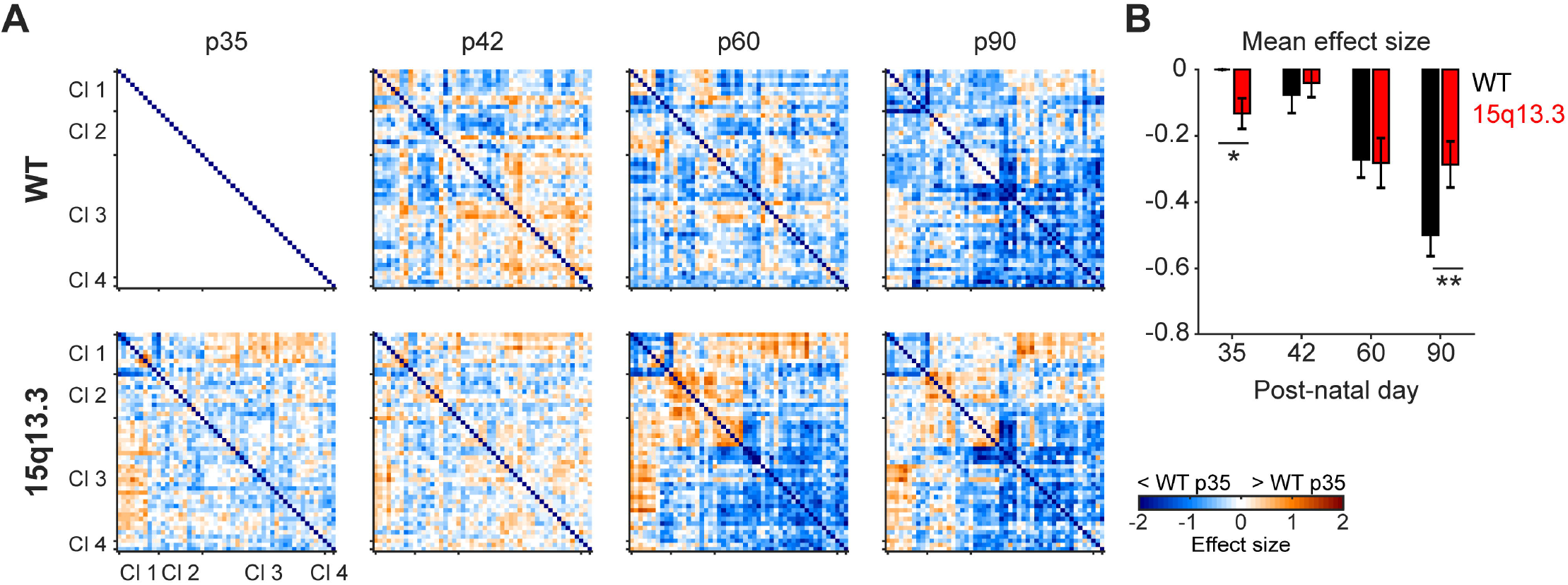
Differential developmental trajectory of functional connectivity in 15q13.3 ^+/−^ mice. (A) Matrices showing the effect size analysis for WT (n=11) and 15q13.3^+/−^ (n=12) mice across multiple time points through development. (B) Mean effect size across multiple developmental stages for WT and 15q13.3^+/−^ mice. 2-way repeated measures ANOVA with Bonferroni’s multiple comparisons correction, genotype x time interaction F_1,49_= 29.5; p<0.001. * p < 0.05; ** p < 0.001 vs WT. Error bars represent ± SEM.

Having identified a large-scale signature of functional connectivity during development in both WT and 15q13.3^+/−^ mice, we were then interested in identifying the specific connections that were different between genotypes. Using NBS with a stringent threshold (t-value ≥ 3, Fig. 4A), we identified a significant network of brain regions that show higher functional connectivity in 15q13.3^+/−^ mice than in WT (Fig. 4). Interestingly, this network revealed a hyperconnectivity pattern of primary and secondary somatosensory cortices (Fig. 4, blue circles) with different thalamic nuclei (Fig. 4, yellow circles) including the habenula, the medio dorsal thalamus, and the hippocampus (Fig. 5, orange circles). Simultaneously, there was an increased intra-amygdala connectivity (Fig. 4, green circles).

**Fig. 4.**
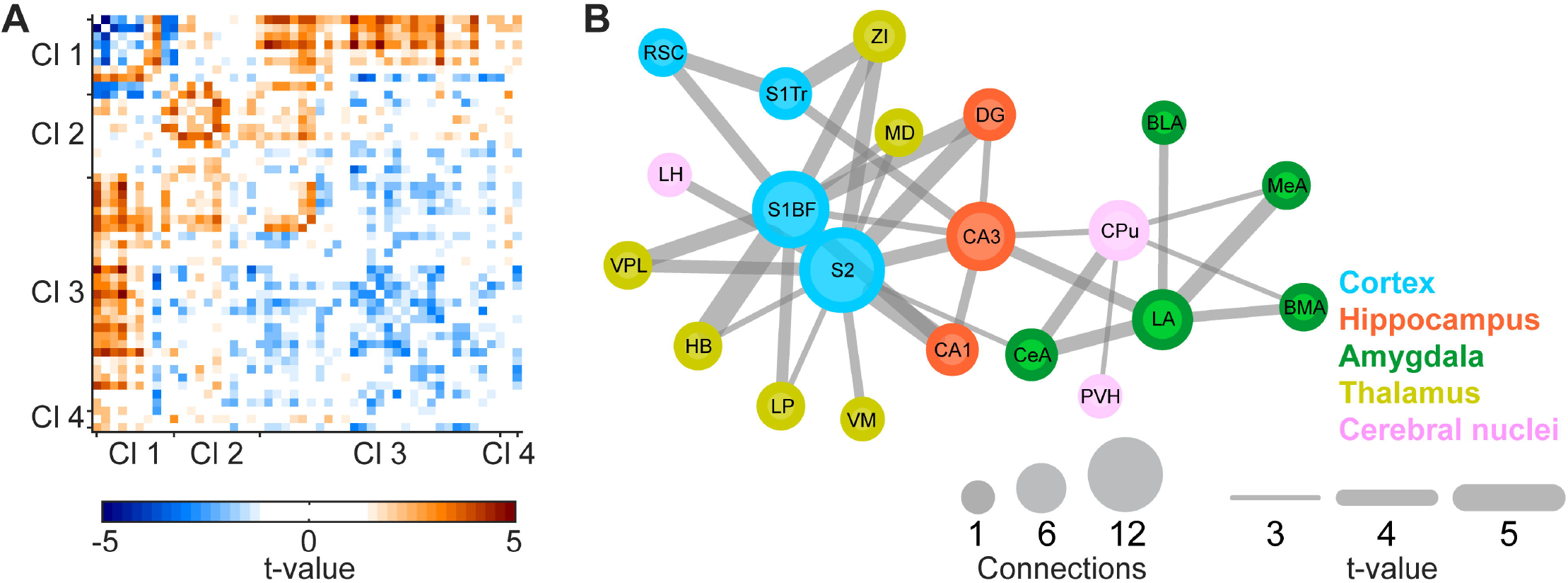
Identification of a large-scale hyper-connected network in 15q13.3 ^+/−^ mice. (A) Scaled matrix of the interregional connections meeting t-value threshold (t-value ≥ 3). (B) Network showing the brain regions (circles) and connections (lines) that were significantly increased in 15q13.3^+/−^ mice with respect to WT. Bilateral regions were grouped for clarity. Abbreviations are defined in Fig. 1.

**Fig. 5.**
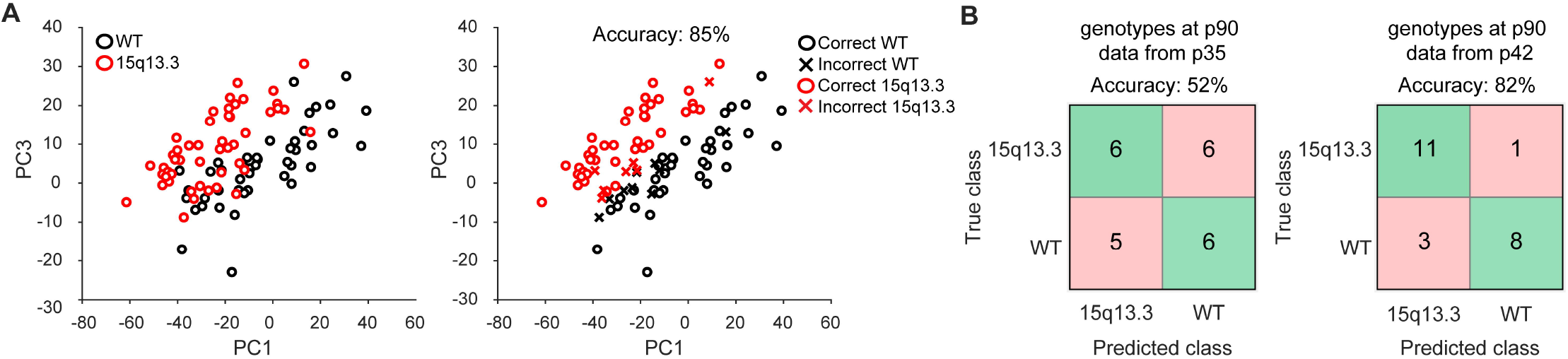
A machine learning algorithm accurately classifies genotype based on functional connectivity. (A) Left, plots showing the loadings for PC1 and PC3 of each individual mouse across all fUS imaging time points. Right, decoding accuracy of the SVM model using the entire dataset, where incorrect predictions are denoted with an x. (B) Decoding accuracy of SVM models trained using functional connectivity data from p35 (left) and p42 (right). Pink boxes denote incorrect classifications and green boxes display correct classifications. WT (n=11) and 15q13.3^+/−^ (n=12).

Since we found that functional connectivity was different in 15q13.3^+/−^ mice, we reasoned that we could use these differences for genotype prediction at later time points. To test this idea, first, we reduced the dimensionality of the entire dataset with PCA and used the loadings of the first three components to train a linear support vector machine (SVM). We found that the first three PCs explained ~37% of the variance of the entire dataset and that the loading values of the individual mice with the same genotype tended to cluster together (Fig. 5A, left panel). Consequently, the SVM model could predict the correct genotypes with 85% accuracy (Fig. 5A, right panel), which was not surprising given that the dataset also included functional connectivity values at p90. Next, we asked whether it was possible to predict the genotypes at p90 by using the functional connectivity values from a distinct time point in development. When we trained the SVM using only functional connectivity data from p35, we found that we could predict the correct genotypes with only 52% accuracy (Fig. 5B, left panel). Interestingly, when we used the functional connectivity from p42, the accuracy to predict the correct genotypes at p90 increased to 82% (Fig. 5B, right panel) which was very similar when used the entire dataset including functional connectivity p90 (Fig. 5A, right panel). Together, these data suggest that there are already changes in functional connectivity at p42 that allow us to predict with high accuracy the genotype later in adulthood.

## Discussion

To the best of our knowledge, this was the first study to monitor functional connectivity through development with fUS imaging and to subsequently use this data for graph theory analysis and prediction modeling. Our network analysis revealed divergent trajectories for 15q13.3^+/−^ mice and WT littermate controls where functional connectivity was reduced for both genotypes with age, but to a lesser extent for 15q13.3^+/−^ mice. We were then able to isolate the distinct differences between genotypes to identify a large-scale network where 15q13.3^+/−^ mice displayed global cortical hyperconnectivity and elevated intra-connectivity within the hippocampus and amygdala. In order to determine the stage of development where the connectivity trajectories bifurcated, we used machine learning to predict genotype classification. We found that the connectivity profile from p42, but not p35, predicted the genotype of individual mice at p90 with the most accuracy. All together, these results suggest a crucial period of network maturation from early to late pubescence that is pivotal in the transition of healthy network connectivity into adulthood.

### Delays and deviations in connectivity patterns in 15q13.3^+/−^ mice through development

When we analyzed the patterns of functional connectivity, we found that in WT mice there was an incremental pattern of decreasing functional connectivity through development (Fig. 3). While there is less data from animal models regarding this phenomenon, these results are in line with longitudinal and cross-sectional studies in humans that show decreases in network functional connectivity from adolescence to adulthood, especially in within network connectivity (Fair et al., 2009; Jolles et al., 2011; Marek et al., 2015; Jiang et al., 2018). These studies found that this was mostly due to decreased segregation and increased integration between networks thought to come about due to synaptic pruning, increased myelination, and an increase in inhibitory signaling (Fair et al., 2007; Supekar et al., 2009). As the brain matures and increases in cognitive functioning, it is necessary to refine network connectivity for faster and more efficient processing. It has been proposed that during development, resting state networks are continually interrogating connections to find the most efficient configuration for a particular function or task (Penn and Shatz, 1999). As these connections are reinforced, either through experience or spontaneous waves of synchrony, redundant connections are pruned leading to an overall refined network that presents with decreased functional connectivity (Fair et al., 2007; Deco et al., 2011), comparable to the trajectory detected in WT mice in this study. It is important to note, however, that the changes in functional connectivity have been presented as differences from WT connectivity on p35. That is, there are still strong functional connections between many of these regions, they are simply reduced in comparison to this time point.

Connectivity in 15q13.3^+/−^ mice, however, displayed a significantly different trajectory of functional connectivity through development. The overall differences in connectivity showed very few changes from p35 to p42. However, the connectivity patterns were noticeably different at p42 in 15q13.3^+/−^ mice compared to WT. At p60 the differences between genotypes were even more evident. There was an overall reduction of functional connectivity in the WT mice and here the 15q13.3^+/−^ mice showed higher magnitudes of change. Hyperconnectivity was evident in global cortical, intra-amygdalar, and intra hippocampal ROIs. At the later adult stage of p90, WT mice showed significant reductions in connectivity, whereas the 15q13.3^+/−^ mice showed few differences from the young adult p60 time point. These results are in line with previous reports showing an increased functional connectivity profile in adult 15q13.3^+/−^ mice (Gass et al., 2016). Furthermore, a reversal of this hyperconnectivity was accomplished through administration of a positive allosteric modulator of the acetylcholine alpha7 receptor, one of the genes in the 15q13.3 locus. The same group later reported increased brain volume (Reinwald et al., 2020), which could be related to reduced pruning compared to WT and reflect the hyperconnectivity signature that we have detected in this study.

### The hyperconnectivity network of 15q13.3^+/−^ mice

A hyperconnected network was revealed using network-based statistics on the entire data set (Fig. 4). In 15q13.3^+/−^ mice, we found hyperconnectivity throughout the primary and secondary somatosensory cortices along with thalamic nuclei and subdivisions of the hippocampus. Thalamocortical hyperconnectivity is frequently reported in patients with first-episode psychosis as well as those with high risk and first-degree relatives (Klingner et al., 2014; Walther et al., 2017; Cao et al., 2018). This aberrant connectivity can have several implications in patients. Firstly, both hypo and hyperkinesia are frequently reported in schizophrenia affected individuals (Lund et al., 1991; Peralta et al., 2010). These features have been associated with differences in myelination and cortical thickness (Bai et al., 2009; Walther et al., 2011), phenomena that are congruent with hyperconnectivity. Secondly, abnormalities in the thalamic relay of sensory information can have implications for the interpretation of external information. Indeed, thalamocortical connectivity has been found to be inversely related to processing speed and attention (Chen et al., 2019) which could be related to abnormal filtering and transmission of sensory information.

We also detected a hyperconnected network within the sub anatomical regions of the amygdala. This feature, in particular, has been linked to some of the positive symptoms of schizophrenia, such as paranoia (Fan et al., 2021; Walther et al., 2021). Increased connectivity of the amygdala with the visual cortices, for example, is prevalent in schizophrenia patients that experience visual hallucinations (Ford et al., 2015). Patients with paranoid ideation have also shown increased connectivity between the amygdala and prefrontal cortex (Fan et al., 2021), a distinguishing trait of schizophrenia and potential diagnostic marker.

We also found increased functional connectivity between hippocampal subregions with amygdala and cortex. Hippocampal hyperactivity is a prominent feature of many psychiatric diseases and especially of schizophrenia. Interestingly, hippocampal connectivity has been correlated to both positive and negative symptoms (Schobel et al., 2009; Duan et al., 2015; Uscătescu et al., 2021). Hallucinations themselves are thought to be in part attributed to the inappropriate recall of memories. Increased resting state connectivity between the hippocampus and somatosensory cortices, even in the absence of a task, can correlate with the severity of hallucination in patients with chronic hallucinations (Sommer et al., 2012). Furthermore, patients with paranoia show increased resting state connectivity between the hippocampus and amygdala, another prominent feature in our study (Walther et al., 2021). Overall, the pattern of functional connectivity in 15q13.3^+/−^ mice seem to recapitulate changes in the pattern of functional connectivity in patients with psychiatric diseases which all hold the potential for biomarkers for schizophrenia diagnosis as well as therapeutic efficacy evaluation.

### Genotype prediction from developmental network signatures

Finally, we used a PCA and machine learning to see if we could accurately predict genotypes using functional connectivity measures from different points in development. We found that the connectivity profile from p35 could only predict the genotype at p90 with 52% accuracy but that data from p42 increased accuracy to 82% (Fig. 5). The ability to predict the conversion from risk to psychosis is of great interest in the treatment and development of therapeutics for schizophrenia. Many of the features detected in this study have been used for that purpose. For example, thalamocortical and amygdalar hyperconnectivity is more prevalent in converters than non-converters (Gee et al., 2012; Ramsay, 2019). Furthermore, the degree of connectivity can be used to predict the time to conversion (Cao et al., 2018). A similar result has been found in the hippocampus, where CA1 hyperactivity could be used to time hippocampal atrophy (Schobel et al., 2013), a feature commonly associated with the conversion to psychosis.

Connectivity profiles can also predict the severity of symptoms (Gheiratmand et al., 2017; Kottaram et al., 2020; Uscătescu et al., 2021), suggesting a direct link between functional connectivity and phenotype. Indeed, measures of connectivity have been shown to most accurately predict symptom severity during follow up after treatment (Collin et al., 2020a). Identification of individuals that are at a high clinical risk for conversion to psychosis in addition to predicting which of these patients will convert, and when that change might occur, could help to thwart illness and improve outcomes. In fact, only a third of subjects deemed clinical high risk eventually receive a formal diagnosis of psychosis (Ellis et al., 2020). Our results show that these same analyses can be applied to a preclinical model, providing a translational tool that may further enhance drug development.

## Conclusion

We have shown that fUS allows for the longitudinal assessment of functional connectivity in mice and that the 15q13.3^+/−^ mouse model can be effectively used to investigate alterations in the trajectory of brain connectivity throughout development. Moreover, we uncovered brain-wide alterations in 15q13.3^+/−^ mice and identified a unique network of aberrant functional hyperconnectivity that included cortical, hippocampal, and amygdalar regions. More importantly, the alterations in functional connectivity observed in the 15q13.3^+/−^ mice resembled those observed in psychiatric patients validating the translational relevance of the mouse model. Finally, we showed that using longitudinal fUS imaging we could identify a connectivity profile in early adolescence that best predicts the network signature during adulthood. Together, these results show that longitudinal fUS imaging in relevant mouse models is a promising approach to improve the development of therapeutic options for psychiatric diseases.

## Acknowledgements

The authors would like to thank Marzieh Funk and Volker Mack for their initial input in the design of the experiments.

## Author details

^1^CNS Discovery Research, Boehringer Ingelheim Pharma GmbH & Co. KG, Biberach an der Riss, Germany.

^2^Institute of Anatomy and Cell Biology, University of Ulm, Ulm, Germany.

## Contributions

G.G-M.: Conceptualization, data collection, data analysis, manuscript writing/editing, final approval of the manuscript. B.H.: Conceptualization, funding acquisition, manuscript editing/review, final approval of the manuscript. H.C.S.: Conceptualization, data analysis, manuscript writing/editing, final approval of the manuscript.

